# Dominant Remodeling of Cattle Rumen Microbiome by *Schedonorus arundinaceus* (Tall Fescue) KY-31 Carrying a Fungal Endophyte

**DOI:** 10.1101/2020.12.18.423411

**Authors:** Bela Haifa Khairunisa, Dwi Susanti, Usha Loganathan, Christopher D. Teutsch, Brian T. Campbell, David Fiske, Carol A. Wilkinson, Frank O. Aylward, Biswarup Mukhopadhyay

## Abstract

Tall fescue KY-31 feeds ~20% of the beef cattle in the United States. It carries a fungal endophyte that produces ergovaline, which causes toxicosis in cattle, leading to $2 billion revenue loss annually. The MaxQ cultivar of the grass is non-toxic, but less attractive economically. To develop ways of mitigating the toxicity, the rumen microbiome of cattle consuming KY-31 and MaxQ have been analyzed, principally for identifying ergovaline transforming microorganisms and often using fecal microbiome as a surrogate. We have hypothesized that KY-31 not only causes toxicosis, but also impacts rumen metabolism broadly, and tested the hypothesis by analyzing rumen microbiome compositions of cattle that grazed MaxQ with an intervening KY-31 grazing period with 16S rRNA-V4 element as identifier. We found that KY-31 remodeled the cellulolytic and saccharolytic communities substantially. This effect was not evident at whole microbiome levels but in the compositions of sessile and planktonic fractions. A move from MaxQ to KY-31 lowered the Firmicutes abundance in the sessile fraction and increased it in planktonic part and caused an opposite effect for Bacteroidetes, although the total abundances of these dominant rumen organisms remained unchanged. In the sessile fraction, the abundances of *Fibrobacter*, which degrades less degradable fibers, and certain cellulolytic Firmicutes such as *Pseudobutyrivibrio* and *Butyrivibrio* 2, dropped, and these losses were apparently compensated by increased occurrences of *Eubacterium* and specific *Ruminococcaceae* and *Lachnospiraceae*. In planktonic fraction the Tenericutes’ abundance increased as saccharolytic Bacteroidetes’ level dropped. Several potential ergovaline degraders were enriched. A return to MaxQ restored the original Firmicutes and Bacteroidetes distributions. However, the *Fibrobacter* and *Butyrivibrio* 2 abundances remained low and their substitutes maintained significant presence. The rumen microbiome was influenced minimally by animals’ fescue toxicosis and was distinct from previously reported fecal microbiomes in composition. In summary, KY-31 and MaxQ cultivars of tall fescue were digested in the cattle rumen with distinct consortia and the KY-31-specific features were dominant. The study highlighted the importance of analyzing sessile and planktonic fractions separately.

## Introduction

In the foregut or rumen of ruminants such as cattle, a complex microbial community anaerobically degrades the feed, generating volatile fatty acids as carbon and energy nutrition for the animals (1). The process also makes livestock major emitters of methane, a potent greenhouse gas (1). For near 100 years there has been intense research focused on rumen microbiome metabolism because of this importance, helping not only to optimize the feed utilization efficiencies in economically important ruminants but also to develop microbial processes for bioenergy production (1–3). In this backdrop, a lack of detailed knowledge of rumen microbiome metabolism in cattle raised on tall fescue is a major gap as it impedes the efforts to utilize this economic grass efficiently for beef and dairy production, to assess the associated impacts on methane emission and for mining this source for novel biocatalysts.

Tall fescue (*Schedonorus arundinaceus* (Schreb.) Dumort., nom. cons. tall fescue) is one of the primary perennial cool season forages that feeds up to 20% of beef cattle in the United States (4). It is grown on more than 35 million acres in the transition zone of Southeastern US, known as the fescue belt (5). In Argentina, Uruguay, and Australia, this forage is grown on over 8.65, 1.24 and 2.72 million acres of the pastures, respectively (6–8). The most widely used variety of the grass is KY-31 and it is popular for its resilience towards pests and poor environmental conditions (5). The resilience of KY-31 is due in part to its symbiotic interaction with a fungal endophyte, *Epichloë coenophiala* (9). The fungus secretes a variety of ergot alkaloids that are beneficial to the grass (9). However, one of these compounds, ergovaline, is toxic to the animals causing Tall Fescue Toxicosis syndrome (10), which results in $2 billion of annual revenue loss for the US beef and dairy industries (11, 12). The use of engineered varieties of tall fescue that either carry a *E. coenophiala* strain that does not produce ergovaline (MaxQ) or are free of the endophyte (KY32, Cajun I, and Bronson) have been promoted as a solution to this problem (9). However, these varieties are less attractive economically due to higher costs for the seeds and pasture management, and KY-31 remains widely used in the fescue belt (13, 14). The selection of fescue toxicosis resistant or tolerant cattle remains an underdeveloped option (5, 15). All these efforts do not consider the possibility that the rumen microbes instigate and could mitigate the above-mentioned toxicity, although there are indications of such possibilities. *In vitro* studies suggest that certain rumen microorganisms transform ergovaline to lysergic acid (16) and it is thought that lysergic acid enters the blood stream of the cattle causing fescue toxicosis (16). The addition of isoflavone producing grasses to a tall fescue diet reduces the severity of fescue toxicosis, and it is thought that isoflavones suppresses ergovaline transforming microbes (11); the mechanism of this suppression is unknown. In search of a microbial basis for the promotion and/or suppression of KY-31 toxicity, the characteristics of the fecal microbiomes of cattle raised on KY-31 and a non-toxic tall fescue have been compared, resulting in the identification of candidate microorganisms (17, 18). Recently, the rumen microbiome of pregnant ewes grazing tall fescue with high and moderate endophyte infection levels in parallel, were analyzed (19). However, these findings do not truly represent the events occurring in the rumen for the following reasons. The feces and rumen harbor distinct microbiomes, sharing ~30% of the species (17, 20). The study with the ewes involved the analysis of rumen fluid drawn via orogastric tube insertion. and therefore, concerned only the planktonic segment and not the whole microbiome as the sessile microbial population was not targeted. We hypothesized that the differences between the microbial systems that KY-31 and MaxQ enrich in cattle rumen are likely broader in nature, going beyond the transformation of ergovaline and covering the overall degradation process occurring both in the planktonic and sessile fractions. Accordingly, we have analyzed the composition of the sessile and planktonic populations of the rumen microbiome of beef cattle that have alternatively grazed MaxQ and KY-31 pastures; the samples were collected through a cannula and two groups of animals with different levels of sensitivities to fescue toxicosis were studied. This is a rare rumen microbiome study with cattle that grazed a pasture and not fed with stockpiled hay or on a feedlot (21–25). For small farmers in Virginia and elsewhere in the US fescue belt, grazing a pasture is the most prevalent way of raising beef cattle before the animals are moved to feed lot for finishing (26). The results of our study have revealed novel type microbial successions and identified KY-31 as a dominant remodeler of sessile and planktonic microbial populations in the rumen, targeting both the cellulolytic and saccharolytic segments.

## Materials and methods

### Preparation of the animals

Eight Hereford-Angus cross steers (age: 18 months; initial body weight: 1079.88 ± 31.92 pounds 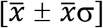 belonging to two groups, sensitive (S, n = 4) and less sensitive (LS, n = 4) to fescue toxicosis, based on the polymorphism in the Dopamine Receptor (DRD2) gene (15) as determined by the T-Snip^TM^ Test (AgBotanica, LLC, Columbia, MO), were used. Each animal was fitted with a 4-inch cannula (Bar Diamond Inc. Parma, ID) at the Virginia-Maryland Veterinary Teaching Hospital (Blacksburg, VA). All of the procedures with the animals were conducted following protocols that were approved by the Virginia Tech Institutional Animal Care and Use Committee (protocol #IACUC 15-065 and 17-171).

### Grazing and rumen sample collection

A rotational grazing study was conducted by moving the animals between two Virginia Tech Agricultural Research and Extension Centers (ARECs) located 130 miles apart. The cattle were first allowed to graze on Jesup MaxQ (27) for 5 months at the Southern Piedmont AREC (SPAREC; Blackstone, VA) (*MaxQ-1 grazing*), then switched to KY-31 for 23 days at the Shenandoah Valley AREC (SVAREC; Raphine, VA) (*KY-31 grazing*), and finally returned to Jesup MaxQ (*MaxQ-2 grazing*). At a point of each grazing period as described in the Results, the samples of rumen contents of each animal were collected from three different locations, top of the mat, middle of the mat, and ventral sac. For the top layer, a sample was taken directly by gloved hand. Samples from the middle of the mat and ventral sac were collected using a custom-made sampling device consisting of a vacuum pump that was connected to a vacuum chamber. Here, each sample was drawn out through the collection tube of the sampling device and deposited directly into the respective sample container. The details of this unit are described in the Supplementary Materials and Methods and Fig S1. Each sample was 500-700 mL in volume and was mixed to get a homogeneity. This large volume allowed better representation of the rumen content in a sample. The liquid and solid fractions of a sample were separated by filtration through a combination of three layers of sterile cheesecloth that had been wrapped separately and baked at 175°C and then transported in sealed 15 ml conical Falcon tubes (catalog number: 62406-200, VWR International, Radnor, PA) on dry ice to the laboratory and stored at −80°C. All containers, tubes and pipes coming into direct contact of the samples were washed prior to using with Tergazyme, a detergent (Alconox Inc, White Plains, NY), and 2% phosphoric acid to remove microbial and nucleic acid contaminations.

### DNA extraction and 16S rDNA amplicon sequencing

From each rumen sample DNA was purified via bead beating and using a modified version of a protocol involving extraction with mixtures of phenol, chloroform, and isoamyl alcohol (25:24:1, v/v/v) and chloroform and isoamyl alcohol (24:1, v/v) (28), isopropanol precipitation and ethanol wash. The details of the method are presented in the Supplementary Materials and Methods. To determine whether a DNA preparation contained inhibitors that could affect downstream processes, near full length 16S rRNA gene was PCR amplified using universal primers 27F and 1525R (29). Metagenomic DNA preparations that yielded the desired amplicons were subjected to paired-end sequencing of the hypervariable region 4 (V4) of the 16S rRNA gene at the Environmental Sample Preparation and Sequencing Facility at the US Department of Energy’s Argonne National Laboratory (ANL). The amplicons were generated using an optimized set of primers, 515F (Parada) – 806R (Apprill), that provides the best coverage of prokaryotic 16S rRNAs (30). The sequencing was performed on the Illumina MiSeq platform (Illumina Inc., San Diego, CA).

### Quantification and statistical analysis of 16S rDNA sequences

Raw sequence data obtained from the ANL were analyzed by the QIIME 2-2019.4 pipeline (31) for preprocessing and removal of contaminants. The Amplicon Sequence Variants (ASVs) were generated by DADA2 (32) and clustered to Operational Taxonomic Units (OTUs) using vsearch (33) at 99% sequence similarity. A pre-trained Naïve Bayes classifier was used to annotate the sequences using the SILVA 132 database (34), and sequences annotated as chloroplast and mitochondria were removed from the dataset. The dataset was imported and analyzed by using the R statistical packages (35).

Alpha diversity metrics were calculated using species richness estimator Chao1 of the microbiomeSeq package (36) with samples rarefied to 5598 sequences per sample (Fig. S2). Pairwise ANOVA (*P* < 0.001) was calculated to determine the significance of the difference between the species richness of two groups. Community comparison between samples was performed using nonmetric multidimensional scaling (NMDS) analysis using Bray-Curtis dissimilarity distances, and unsupervised principal component analysis (PCA) using the data normalized into relative abundance with total sum scaling (TSS) and transformed into its logarithmic values using centered log ratio (CLR) as described (37, 38). Phyloseq (39) and vegan (40) were used in R to calculate sample distances (Bray-Curtis and UniFrac with average as method), perform NMDS analysis, and compute a dendrogram of the hierarchical clustering. The TSS and CLR data transformation, as well as sample ordination on PCA were performed using mixMC (37). A non-parametric permutational analysis of variance (PERMANOVA) and analysis of similarities (ANOSIM) of adonis function in vegan (40) were used to determine which sample parameters (genotype, rumen sample fraction, rumen depths, tall fescue variety, and grazing transition) caused sample clustering (permutation: 999, *P* < 0.05). Relative abundance of the OTUs in various samples were estimated using phyloseq (39), microbiome (41), and DESeq2 (42) packages. Core microbial community in a rumen sample was characterized from TSS normalized data at 95% prevalence threshold across all samples. The assessment of differential abundance of microbial species between two grazing periods with respect to either sample fractions (solid or liquid) or sensitivities to fescue toxicosis was performed using DESeq2 (42) with Wald p-value < 0.001, fitType = “parametric”, and sfType = “poscounts”. Non-parametric Kruskal-Wallis (*P*_Kruskal-Wallis_ *=* 0.05) and Wilcoxon test with continuity correction (*P*_Wilcoxon_ = 0.05) were performed to determine the significance differences in the relative abundance of a microbial species across rumen fraction and grazing transition.

### Data availability

We are in the process of submitting the raw sequences of the 16S rDNA-V4 amplicons to the National Center for Biotechnology Information Sequence Read Archive (NCBI SRA). We will list the accession number for this submission as soon as it becomes available and certainly prior to the publication of the manuscript.

## Results

### Rumen samples and Sequences of 16S rDNA V4 Amplicons

The first set of samples of rumen contents were collected at the end of the initial 5 month-long MaxQ grazing period (MaxQ-1). The animals then grazed KY-31 for 24 days. At the end of this period (KY-31), a second set of samples were collected, and the animals were placed on MaxQ grazing (MaxQ-2). After 23 days of the MaxQ-2 phase, the last set of samples were collected. Under all these grazing conditions, the rumen pH remained constant (7.00 ± 0.24); the same was the case for the animal’s rectal temperature and average daily gain. Sampling in triplicate from three rumen locations, top and middle of the mat and the ventral sac, of eight animals and for three grazing periods produced 216 solid and 216 liquid fractions. Sequencing of the 16S rDNA-V4 elements of the DNA preparations generated from these samples provided 6.2 million sequences with 9,403 ASVs. A clustering of the data at the 99% similarity threshold produced 5,547 unique OTUs. Further filtering, including the removal of 12 samples with less than 5,200 reads to avoid introducing noise (Table S1), provided a final data set comprised of 5,488 OTUs from 420 samples.

### Alpha and Beta Diversity of Tall Fescue Rumen Microbiome

Overall, the species richness in the microbiome of the solid fractions of the rumen contents was similar to that of the respective liquid fraction (Fig. 1). In terms of the location within the rumen, the top of the mat harbored a less diverse microbial population than the middle of the mat and the ventral sac (Fig. 1). The grazing transitions altered the species richness significantly (Fig. 1), distinguishing the MaxQ-2 set (return from KY-31 to MaxQ) with higher species richness than MaxQ-1 and KY-31 (Fig. 1); the richness was the lowest under KY-31 grazing (Fig. 1). For both MaxQ grazing periods (MaxQ-1 and MaxQ-2), the solid fraction presented a microbiome with higher species richness than the respective liquid fraction, and the situation was opposite with KY-31 (Fig.1). The KY-31 grazing resulted into a less diverse sessile microbial community (Fig. 1). The rumen microbiome during the MaxQ-2 grazing of cattle that were less sensitive to fescue toxicosis had a slightly higher species richness compared to rest five sets (Fig. 1).

**Figure 1.**
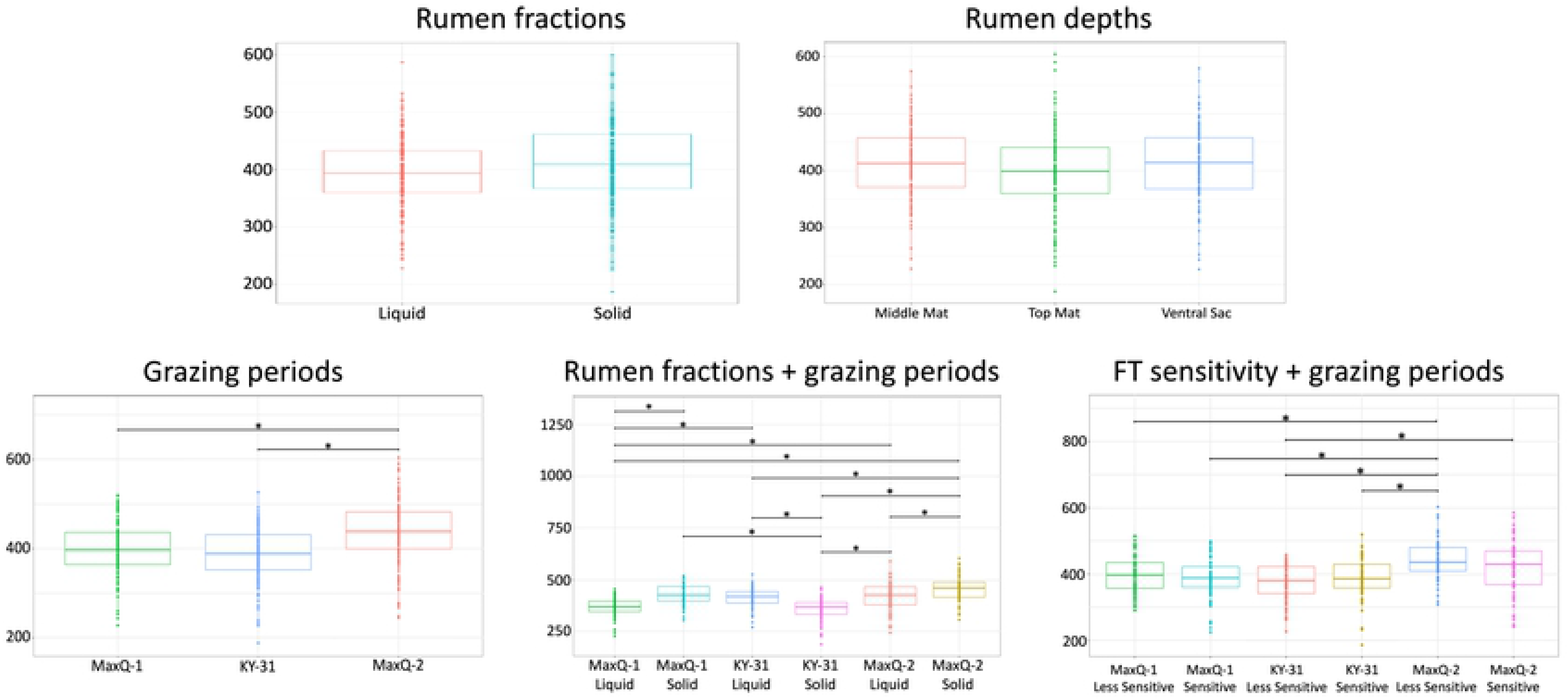
Species richness of rumen microbiome of cattle grazing tall fescue. The cattle grazed the following tall fescue pastures in the sequence shown (designation of the grazing period): Jesup Max Q (MaxQ-1); KY-31 (KY-31); Jesup Max Q (MaxQ-2). Analysis of alpha diversity occurred across rumen fractions (solid and liquid), rumen depths, grazing periods, combination of rumen fractions and grazing periods, and combination of sensitivities to fescue toxicosis and grazing periods. Prior to species richness analysis, all of the samples were rarefied to 5598 reads, which was the lowest number of reads present in a single sample (Fig. S2). Horizontal bars link the samples compared in statistical analyses. *Represent *P* < 0.001 in a pairwise ANOVA statistical test.

Community comparison via Bray-Curtis and Euclidean distances ordination (NMDS and PCA, respectively) showed clear separations between the solid (sessile) and liquid (planktonic) fractions of the microbiomes and the tall fescue types used (Fig. 2A-B). The distinctions between the rumen microbiomes on MaxQ-1, KY-31, and MaxQ-2 were more pronounced for the liquid fractions, and here MaxQ-1 samples were clearly separated (Fig. 2A-B). A similar separation, although to a lesser extent, was observed for the solid fractions. Interestingly, MaxQ-2 had some commonalities with KY-31 but not MaxQ-1 (Fig. 2A-B).

**Figure 2.**
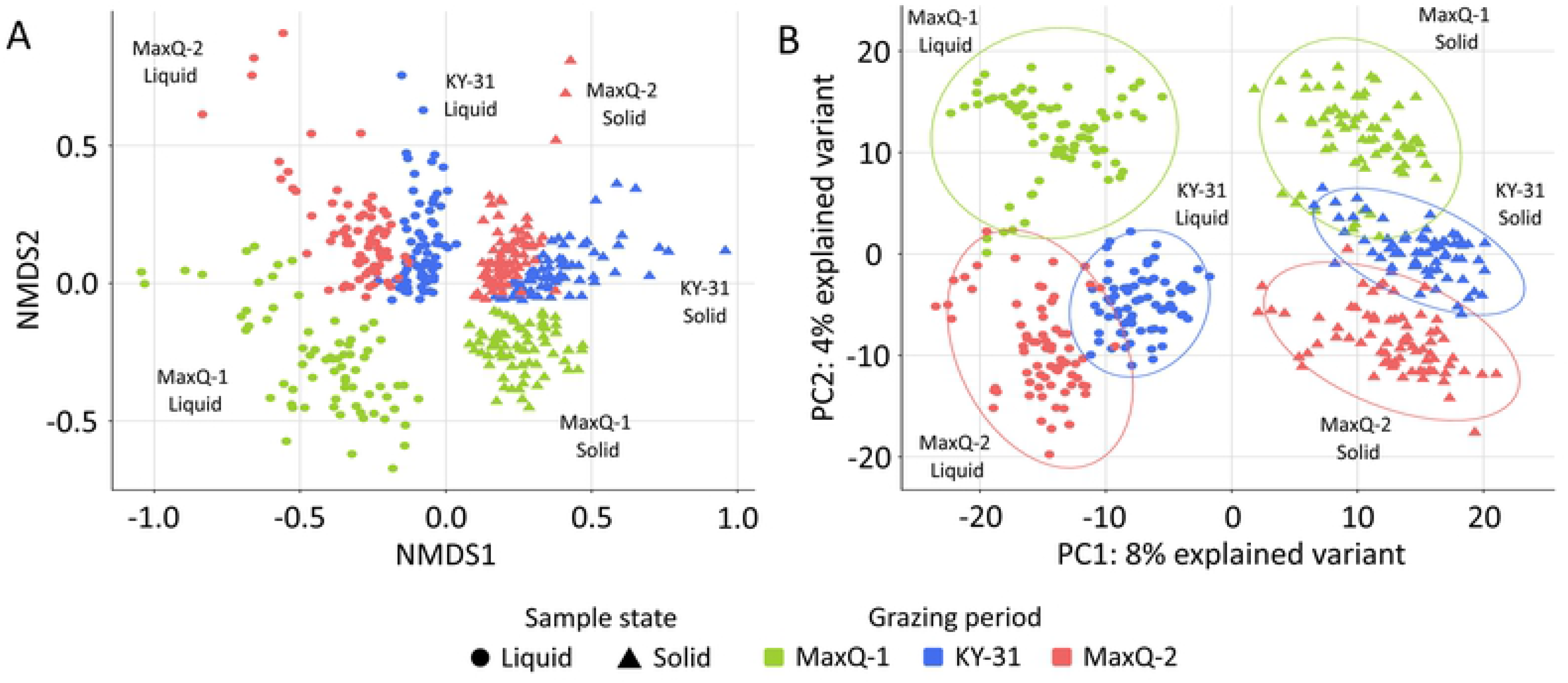
Comparison of the rumen microbiome composition of cattle grazing Max-Q and KY-31 tall fescue pasture. A non-metric multidimensional scaling (NMDS) ordination of the Bray-Curtis dissimilarity distances of tall fescue rumen microbiome **(A)** and PCA of the TSS normalized and CLR transformed rumen microbiome data **(B).** The rumen microbiome was clustered based on the rumen fraction and grazing transition.

Quantitative assessment of the parameters that caused sample clustering was performed using permutational multivariate analysis of variance (PERMANOVA; 999 permutation and P_PERMANOVA_ = 1e^−04^) and similarities (ANOSIM; P_ANOSIM_ = 0.001) of the Bray-Curtis dissimilarity distance. Table 1 presents the *P*-values obtained from these analyses. The results show that the rumen depth was an insignificant parameter in both of these tests (P_PERMANOVA_ = 0.3045 and P_ANOSIM_ = 0.104), indicating that it did not contribute to the separation of the samples. In both PERMANOVA and ANOSIM analysis, the animal genotypes were found to be a significant parameter (P_PERMANOVA_ = 1e^−04^ and P_ANOSIM_ = 0.001) (Table 1), although it did not cause separation on the ordination plot (Fig. 3).

**Table 1.** PERMANOVA and ANOSIM statistical analysis of the sample parameters.

| Statistical Method | Parameter | p-value | R value |
| --- | --- | --- | --- |
| **PERMANOVA** | Tall Fescue variety | 1e-04* | 0.04835 |
|  | Genotypic factor | 1e-04* | 0.08395 |
|  | Rumen fraction | 1e-04* | 0.13672 |
|  | Rumen depth | 0.3045 | 0.00508 |
|  | Grazing transition and rumen fraction | 1e-04* | 0.08488 |
| **ANOSIM** | Tall Fescue variety | 0.001* | 0.06795 |
|  | Genotypic factor | 0.001* | 0.1123 |
|  | Rumen fraction | 0.001* | 0.5885 |
|  | Rumen depth | 0.104 | 0.003578 |
|  | Grazing transition and rumen fraction | 0.001* | 0.6154 |

**Legend:** PERMANOVA and ANOSIM were conducted with 999 permutation and *P*-value below 0.05 was considered significant.

**Figure 3.**
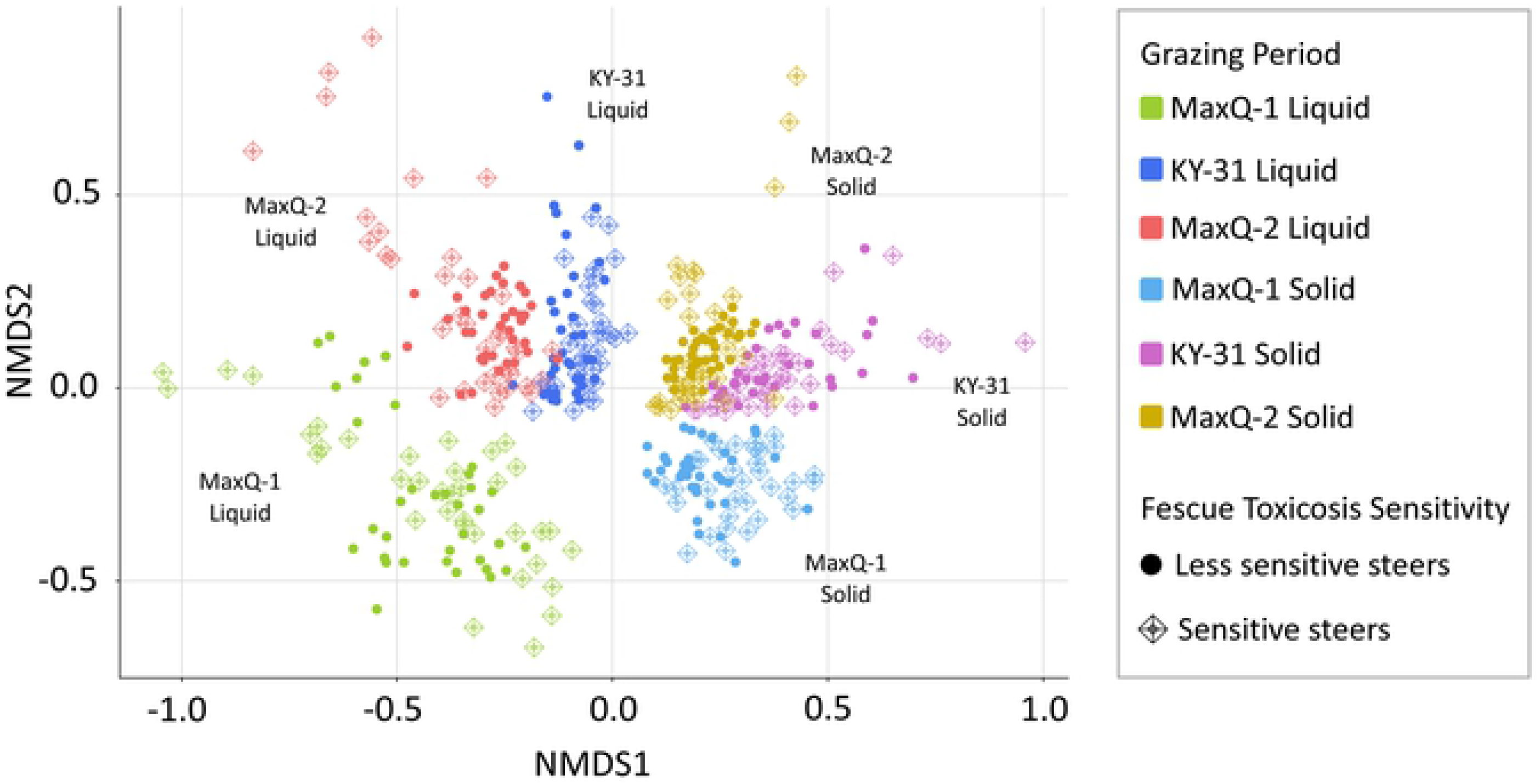
Effect of fescue toxicosis sensitivity of the cattle on their rumen microbiome compositions. The results shown are from an NMDS analysis of the data set based on Bray-Curtis dissimilarity distances.

### Tall fescue cultivar change-driven alterations in the core rumen microbiome

Under all three grazing conditions the rumen core microbiome was represented by 25 OTUs and contained bacteria with the taxonomic identities as shown in Fig. 4B and Table S2. There were remarkable alterations in the locations of the Bacteroidetes, Firmicutes, Fibrobacter, and Lentisphaerae populations within the microbiome due to changes in the tall fescue type (Fig. 4A-B). We describe this effect with respect to the sessile and planktonic populations, separately. Following the transfer from MaxQ-1 to KY-31, there was a significant increase in the number of Bacteroidetes OTUs in the sessile fraction (MaxQ-1, 41.78%; KY-31, 46.44%) and then a decrease during MaxQ-2 grazing (35.33%). For Firmicutes, the observation was the opposite, as in the solid samples the abundance slightly dipped during KY-31 grazing (from 41.47% in MaxQ-1 to 40.38% in KY-31) before rising to 49.11% under MaxQ-2. In the liquid fraction, the abundance of Bacteroidetes OTUs continuously dropped from 41.49% in MaxQ-1 to 38.74% and 36.47% in KY-31 and MaxQ-2, respectively. These losses of Bacteroidetes in the liquid samples paralleled the gains in the abundance of planktonic Firmicutes (41.07%, 43.84%, and 46.63% for MaxQ-1, KY-31, and MaxQ-2, respectively).

**Figure 4.**
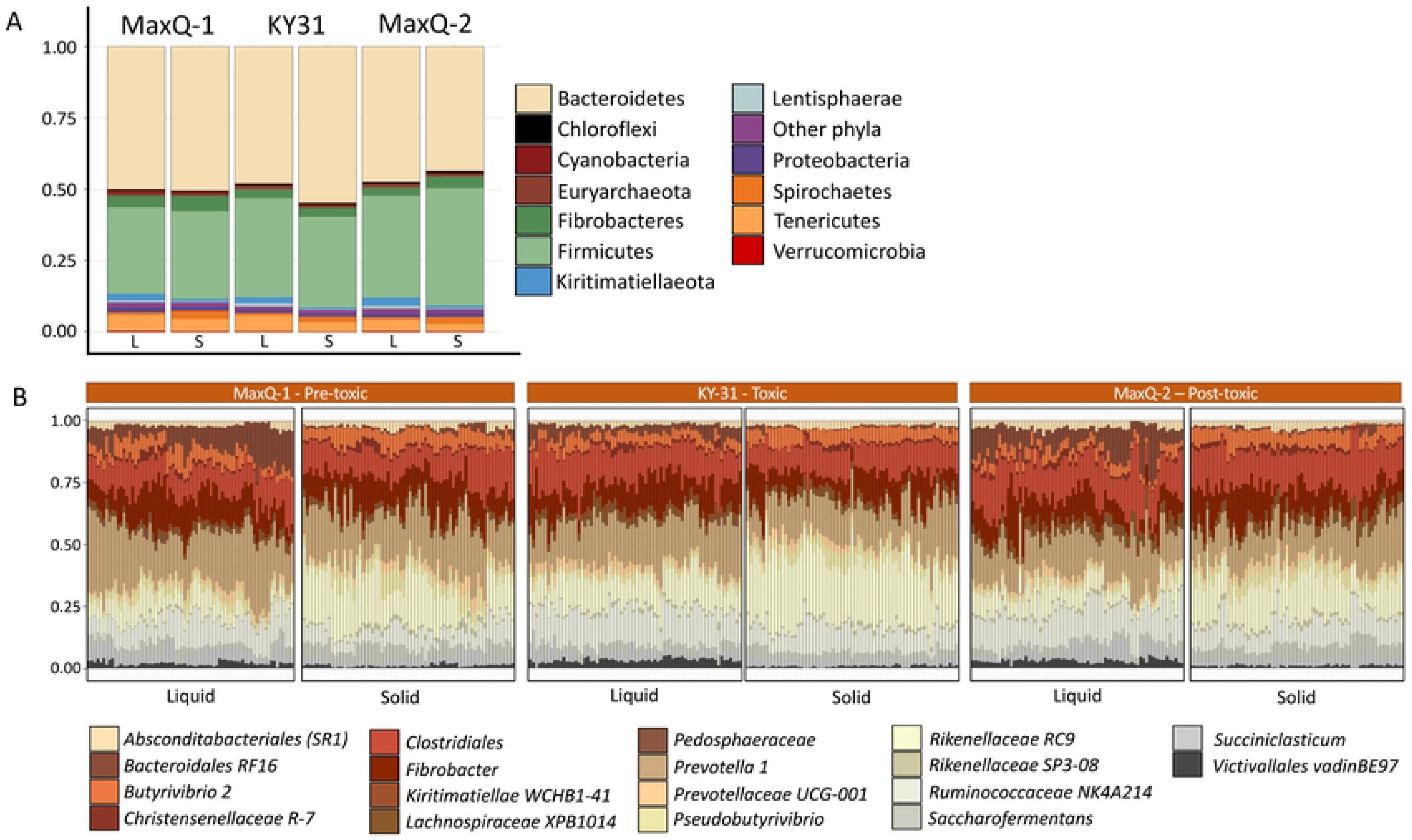
Core microbiome composition shifts in the solid and liquid phases of the rumen content due to grazing transitions. Microbiome composition at the phylum-level **(A)** and core microbiome composition of 25 OTUs that were present at 95% prevalence threshold across all samples **(B)**. Assigned lowest level taxonomic annotations are shown with different colors as listed at the bottom of the figure. Y-axis represent percent of relative abundance.

At 95% sample prevalence, no Euryarchaea OTUs were detected in the core rumen microbiome. The prevalence thresholds that allowed the detection of the Euryarchaeota OTUs were 70%, 80%, and 85% for KY-31, MaxQ-1, and MaxQ-2, respectively. At 70% prevalence, the Euryarchaeota was found to be represented by the *Methanobacteriaceae* and *Methanomethylophilaceae* families under all grazing conditions. Further analysis of all the data without a prevalence cutoff showed that methanogen abundance did not change significantly with anyone of the parameters studied (Fig. 5), except the KY-31 grazing slightly enhanced the abundance of species from the *Methanosarcinaceae* family in the liquid fraction (Fig. 5).

**Figure 5.**
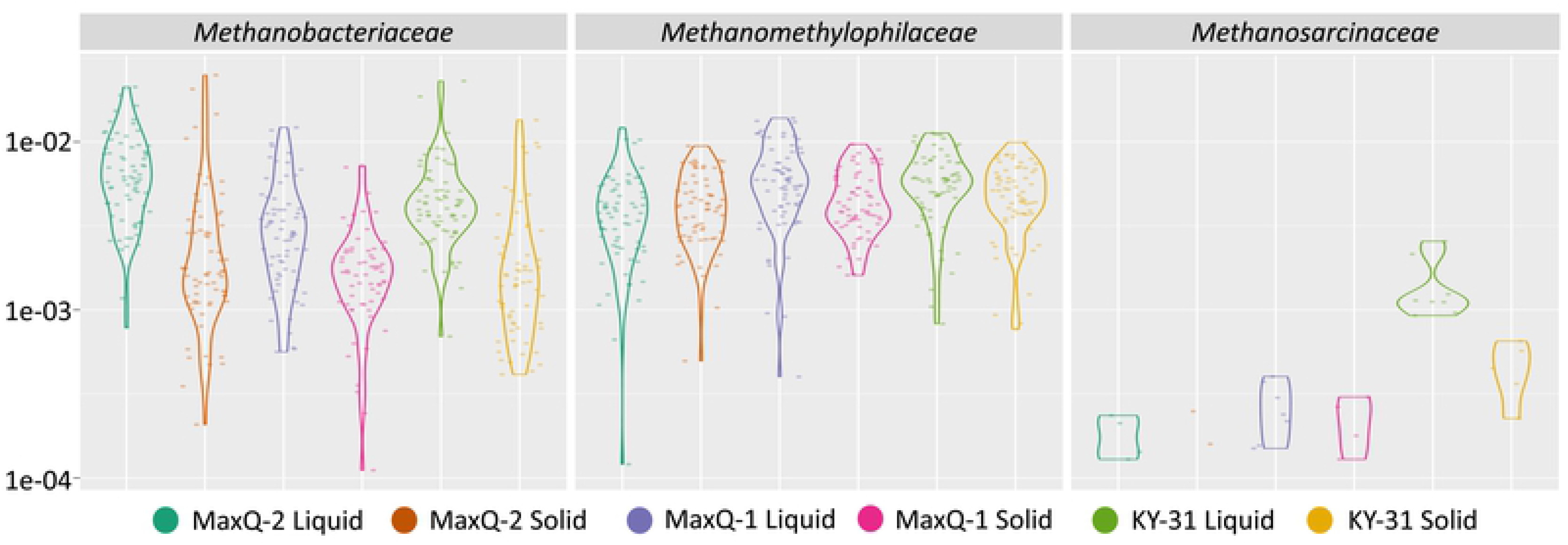
Euryarchaeota abundance in tall fescue rumen microbiome. Y-axis: relative abundance of Euryarchaeota family in tall fescue rumen microbiome. X-axis: rumen fraction in each of the grazing transition are shown with different colors as indicated at the bottom of the figure. The cattle grazed the following tall fescue pastures in the sequence shown (designation of the grazing period): Max-Q (MaxQ-1); KY-31 (KY-31); Max-Q (MaxQ-2). Liquid and Solid: respective phases of rumen contents.

The core bacterial microbiomes of steer grazing MaxQ (pre- and post-KY-31) and KY-31 tall fescue pasture had clear differences (Fig. 4B). *Prevotella 1* genus of the phylum Bacteroidetes had higher abundance in MaxQ-1 (21%) compared to KY-31 (17%) and MaxQ-2 (16%) grazing; the %values are calculated with respect to the core microbiome OTUs. Transition to KY-31 grazing promoted the growth of *RC9* species of the *Rikenellaceae* family (Bacteroidetes phylum) significantly, as the abundance of the respective OTUs increased from 6.47% and 16.41% in the liquid and solid fractions under MaxQ-1, respectively, to 12.15% and 25.71% in the corresponding sets during KY-31 grazing. Compared to the MaxQ-1, the KY-31 grazing provided higher abundances for bacteria belonging to the candidate division vadinBE97 of the Lentisphaerae phylum, and the Firmicutes of *NK4A214* and *XPB1014* genera from the *Ruminococcaceae* and *Lachnospiraceae* families, respectively, and of unclassified Clostridiales order (Fig. 4B). Under KY-31 grazing, the population of *Fibrobacter* species of the Fibrobacteres phylum, and *Succiniclasticum*, and *Butyrivibrio 2* species of the Firmicutes and the *RF16* family of the Bacteroidales order decreased, and for the last group the drop was substantial (about 50%) (Fig. 4B and Table S2).

The transition from KY-31 back to MaxQ resulted in another reconstruction of the core microbiome characteristics. Some of the bacterial species, for which the abundance decreased during KY-31, regained their presence under MaxQ-2 to the level that was seen under MaxQ-1. This was the case for the bacteria of the RF16 family (Bacteroidales order), for which the abundance decreased from 9.74% in MaxQ-1 to 4.22% in KY-31 and rose to 9.41% under MaxQ-2 grazing. Similar was the observation for *Succiniclasticum* species of the Firmicutes phylum (Fig 4B). The abundances of *Pseudobutyrivibrio* and *Butyrivibrio 2* species of the Firmicutes phyla and *Fibrobacter* remained low and did not recover in the MaxQ-2 grazing (Table S2). A non-parametric Kruskal-Wallis and Wilcoxon rank sum test (43–45) also showed a reduction in the abundance of Fibrobacter in sessile fraction due to shift from MaxQ to KY-31 and a poor recovery upon return to MaxQ (Fig. 6).

**Figure 6.**
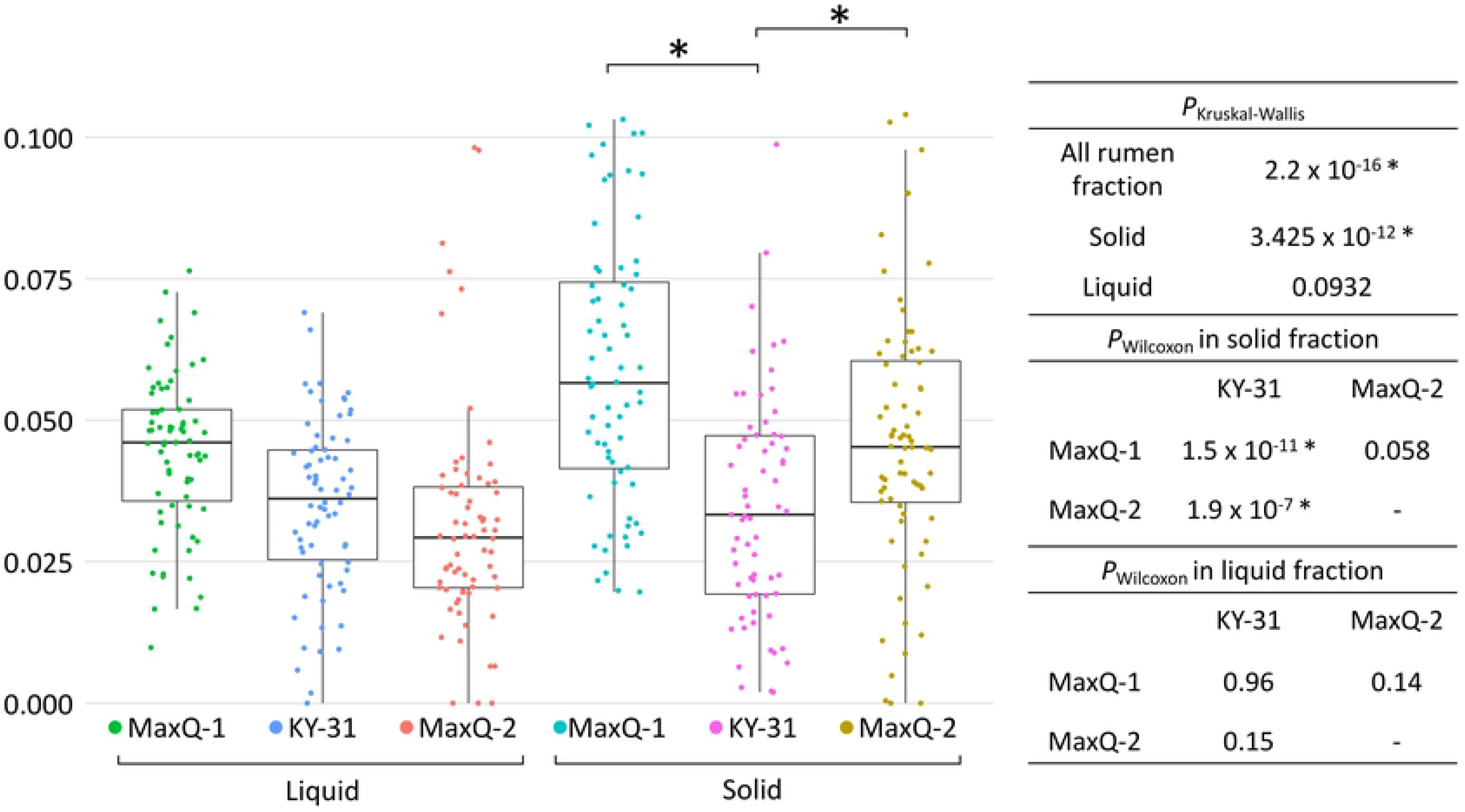
Relative abundance of *Fibrobacter* in the rumen microbiome during tall fescue grazing with respect to grazing periods, MaxQ-1, KY-31, and MaxQ-2, and rumen fractions. Boxplot of the relative abundance is shown on the left figure whereas *P*-values of Kruskal-Wallis (*P*_Kruskal-Wallis_ = 0.05) and pairwise Wilcoxon rank sum test (*P*_Wilcoxon_ = 0.05) with respect to grazing periods and rumen fraction are shown on the chart.

### Differential abundance analysis of grazing transition driven shift in microbiome

We performed a DESeq2 analysis (42) to identify the shifts in the microbiome composition, where enriched OTUs in each grazing comparison (MaxQ-1 vs KY-31; KY-31 vs MaxQ-2; and MaxQ-1 vs MaxQ-2) were defined as those with P_Wald_ less than 0.001 (Table S3). A similar analysis was performed with rumen fractions as a parameter (Table S4). The results of these analysis for MaxQ-1 vs KY-31 and KY-31 vs MaxQ-2 are summarized in Fig 7, both in terms of the total microbiomes as well as sessile and planktonic fractions, and that for MaxQ-1 vs MaxQ-2 appear in Fig 8. The fiber-adherent community in the rumen during MaxQ-1 grazing had four-fold more abundance for *Fibrobacter* than that with KY-31 and similarly higher abundance OTUs were seen for the genus *Saccharofermentans*, family of *Lachnospiraceae AC2044,* Spirochaetes MVP-15 order, and WCHB1-41 class from Kiritimatiellae class (Fig. 7). For the Bacteroidetes phyla, a transfer from MaxQ-1 to KY-31, enriched the rumen microbiome greatly with an uncultured bacterium from the Bacteroidales order and *Rikenellaceae RC9*, *Prevotella* 1, and *Paraprevotella* species (Fig. 7). For the Firmicutes phyla, this grazing transition enriched only one species that belonged to the genus of *Lachnospiraceae FCS020*. After a month from the return to MaxQ (MaxQ-2), the abundance of a set of OTUs was elevated and these had shared annotations but not identical sequences with a group of OTUs in MaxQ-1. This observation could be interpreted as the occurrence of two groups of organisms with close but not exactly the same genetic identities in these two sets. This group was comprised of *Lachnoclostridium 10*, *Christensenellaceae R-7*, *Treponema 2*, *Anaeroplasma, Saccharofermentans*, and *Lachnospiraceae AC2044* genera and uncultured bacteria belonging to the BS11 gut group of Bacteroidales order (Fig. 7).

**Figure 7.**
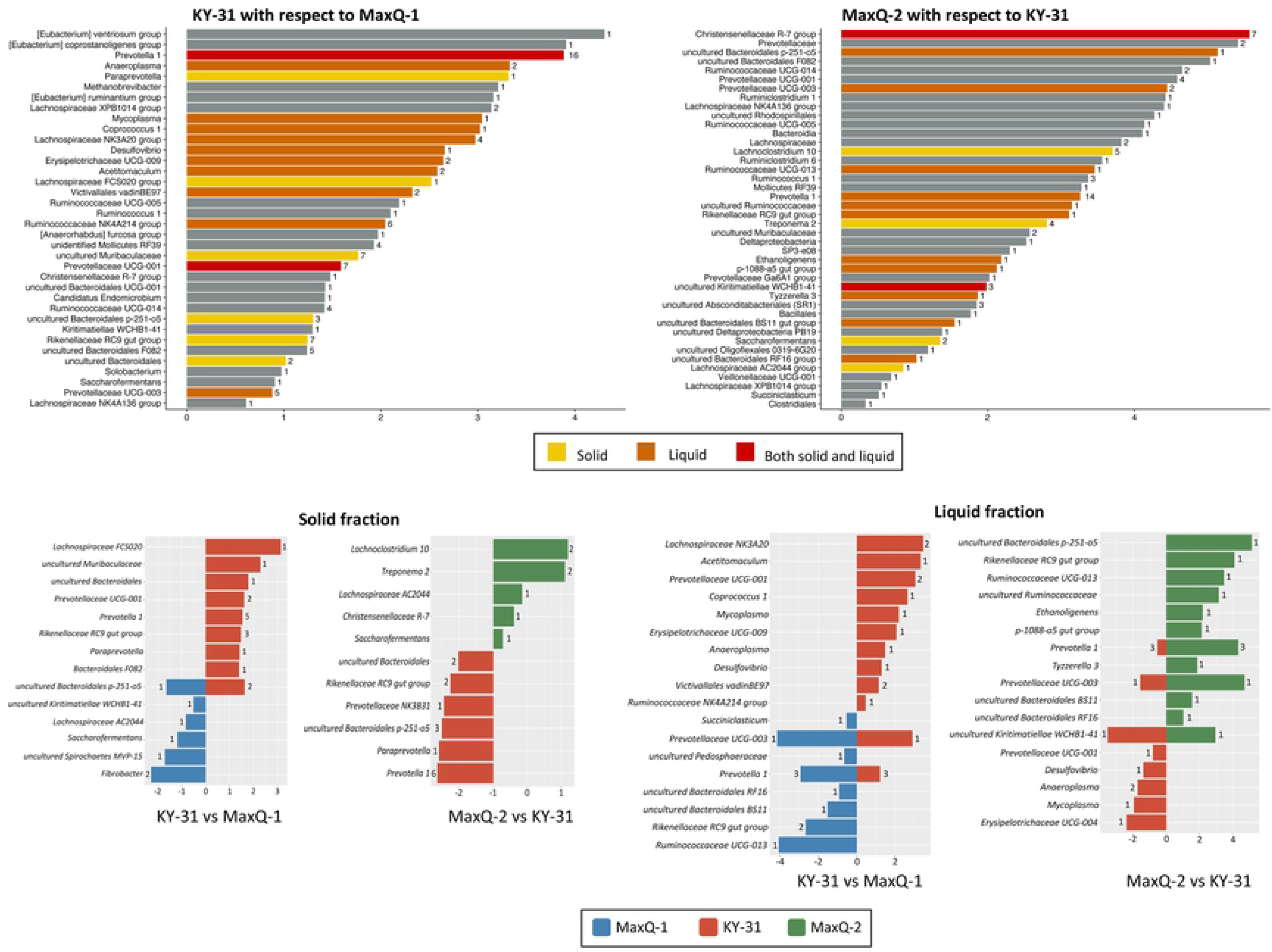
Grazing transition-associated shifts in the compositions of rumen microbiome. **Top plots:** Differential abundances of bacteria in KY-31 with respect to MaxQ-1 and in MaxQ-2 with respect to KY-31. **Bottom plots:** Sessile and planktonic populations for the Top Plots. Average Log2Fold values are shown on the x-axis and the number of OTUs representing an assigned lowest taxonomic annotation is shown on the side of the respective bar.

**Figure 8.**
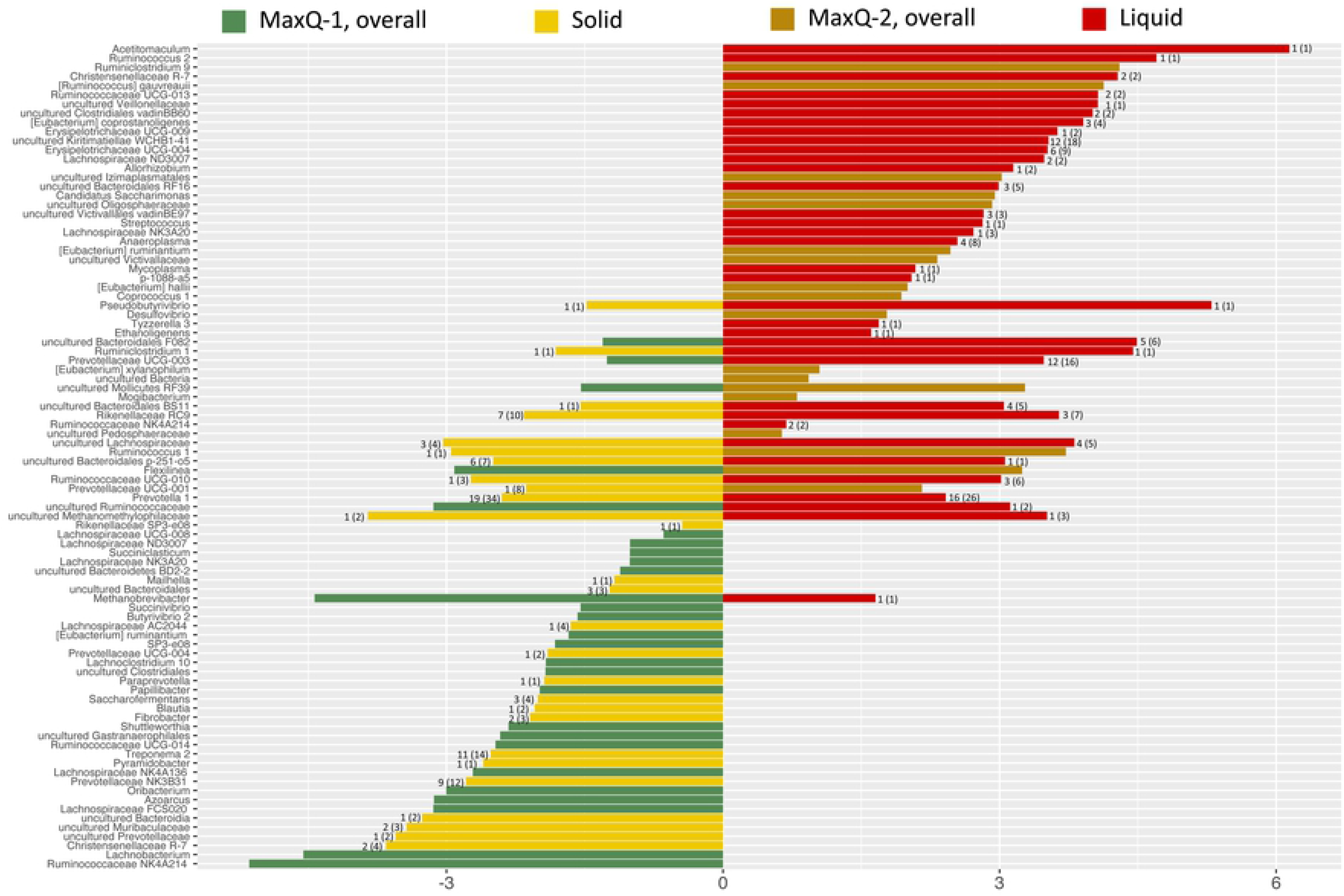
Microbial community comparison between two MaxQ grazing periods. Differentially abundant bacteria in: MaxQ-1 with respect to MaxQ-2; MaxQ-2 with respect to MaxQ-1 Average Log2Fold values are shown on the X-axis and the number of OTUs representing an assigned lowest taxonomic annotation is shown on the side of the respective bar.

The planktonic population in MaxQ-1 microbiome was characterized by high abundances of the following organisms: *Succiniclasticum*, *Shuttleworthia*, *Prevotellaceae UCG-003*, and *Ruminococcaceae UCG-013* genera, and uncultured bacteria from *Pedosphaeraceae* family, and RF16 and BS11 families of Bacteroidales order (Fig. 7). In the liquid fraction-associated microbiome during KY-31 grazing, the abundances of *Anaeroplasma* and *Mycoplasma* species of the Tenericutes phyla, *Desulfovibrio 2* species from the Proteobacteria phyla, organisms from the Firmicutes phyla belonging to the families of *Erysipelotrichaceae* (UCG-004 and UCG-009), *Lachnospiraceae* (*Acetitomaculum* and *Lachnospiraceae* NK3A20 species), and *Ruminococcaceae* NK4A214, WCHB1-41 class of the Kiritimatiellaeota phylum, and Victivallales vadinBE97 order of Lentisphaerae phylum increased (Fig. 7). The return to MaxQ (MaxQ-2) altered the community composition again, as *Rikenellaceae RC9*, *Ruminococcaceae UCG-013*, *Ethanoligenes*, and *Tyzzerella 3* of Firmicutes phylum, *Prevotellaceae UCG-001* of the Bacteroidetes phylum, and *Pirellulaceae p-1088-a5* gut group of Planctomycetes phylum were found with significantly higher abundance here.

Of the two MaxQ grazing periods, MaxQ-1 provided a higher abundance of Bacteroidetes members such as *Paraprevotella*, *Saccharofermentans*, and uncultured species of the *Muribaculaceae* family, the genera of *Blautia, Ruminococcus 1*, and *Lachnospiraceae AC2044* genera of the Firmicutes phylum, cellulolytic *Fibrobacter* from the Fibrobacteres phylum, and a *Synergistaceae* family member *Pyramidobacter* in the cattle of rumen (Fig. 8. Bacteria of the *Treponema 2* genus from the Spirochaetes phylum as well as those from the *Prevotellaceae* family, *NK3B31*, were found with a relatively exceptionally high OTU numbers (11 and 9, respectively) during this grazing (Fig. 8). In the MaxQ-2 grazing, *Victivallales vadinBE97* from Lentisphaera phylum, a Planctomycetes member *Pirellulaceae p-1088-a5* gut group, *Anaeroplasma* and *Mycoplasma* species from the Tenericutes phylum, *Pseudobutyrivibrio, Erysipelotrichaceae* members *UCG-009* and *UCG-004*, *Lachnospiraceae NK3A20* and *ND3007*, *Acetitomaculum* and *Tyzzerella 3* species from the Firmicutes phyla, and a *Methanobrevibacter* species of the *Methanobacteriaceae* family from Euryarchaeota phylum, were found to be enriched in this grazing (Fig. 8). Bacteria of the WCHB1-41 class from the Kiritimatiellaeota phylum, showed the highest OTU numbers (12 OTUs) while three Firmicutes genera, *Ruminococcus 2*, *Pseudobutyrivobrio*, and a known rumen acetogen *Acetitomaculum* displayed the most significant abundances (26, 40, and 70-fold higher, respectively) in the MaxQ-2 grazing (Fig. 8). An interesting pattern was seen for *Prevotella 1* species from Bacteroidetes phyla, as a total of 60 distinct OTUs for this genus were detected in the samples from MaxQ-1 and MaxQ-2 periods. While 34 *Prevotella 1* OTUs were found during MaxQ-1 grazing of which 19 were highly abundant, the corresponding values were 26 and 16 in the MaxQ-2 period, respectively. Another significant observation was that certain *Eubacterium* species, which are Firmicutes, were abundant during the KY-31 grazing irrespective of the rumen fractions (Fig. 7) and *Eubacterium ruminantium* in particular was present at a high abundance in the MaxQ-2 period (Fig 8).

### Host genotype-associated microbial community characteristics

Bacteroidetes OTUs were significantly more abundant in the less sensitive steers (Fig. 9 and Table S5) and also higher number of OTUs from the Proteobacteria phyla such as those corresponding to the *Sutterella* genus and Rhodospirillales and Deltaproteobacteria PB19 orders were also associated with this host genotype. In the rumen of this group the OTUs that had higher abundance during MaxQ-1 and MaxQ-2 grazing but were less so in the KY-31 grazing mostly belonged to *Prevotellaceae*, *Marinilabiliceae*, and *Rikenellaceae* families, and some of these were annotated as unclassified Rhodospirillales order from Proteobacteria phylum and Gastranaerophilales from the newly defined phylum of Melainabacteria that is closed-related to Cyanobacteria. The cellulolytic *Fibrobacter* species were 22- and 59-fold more abundant in the less sensitive steers during MaxQ-1 and MaxQ-2 grazing, respectively (Fig. 9), and such a distinction was not seen during KY-31 grazing, where the abundances of these organisms were generally low.

**Figure 9.**
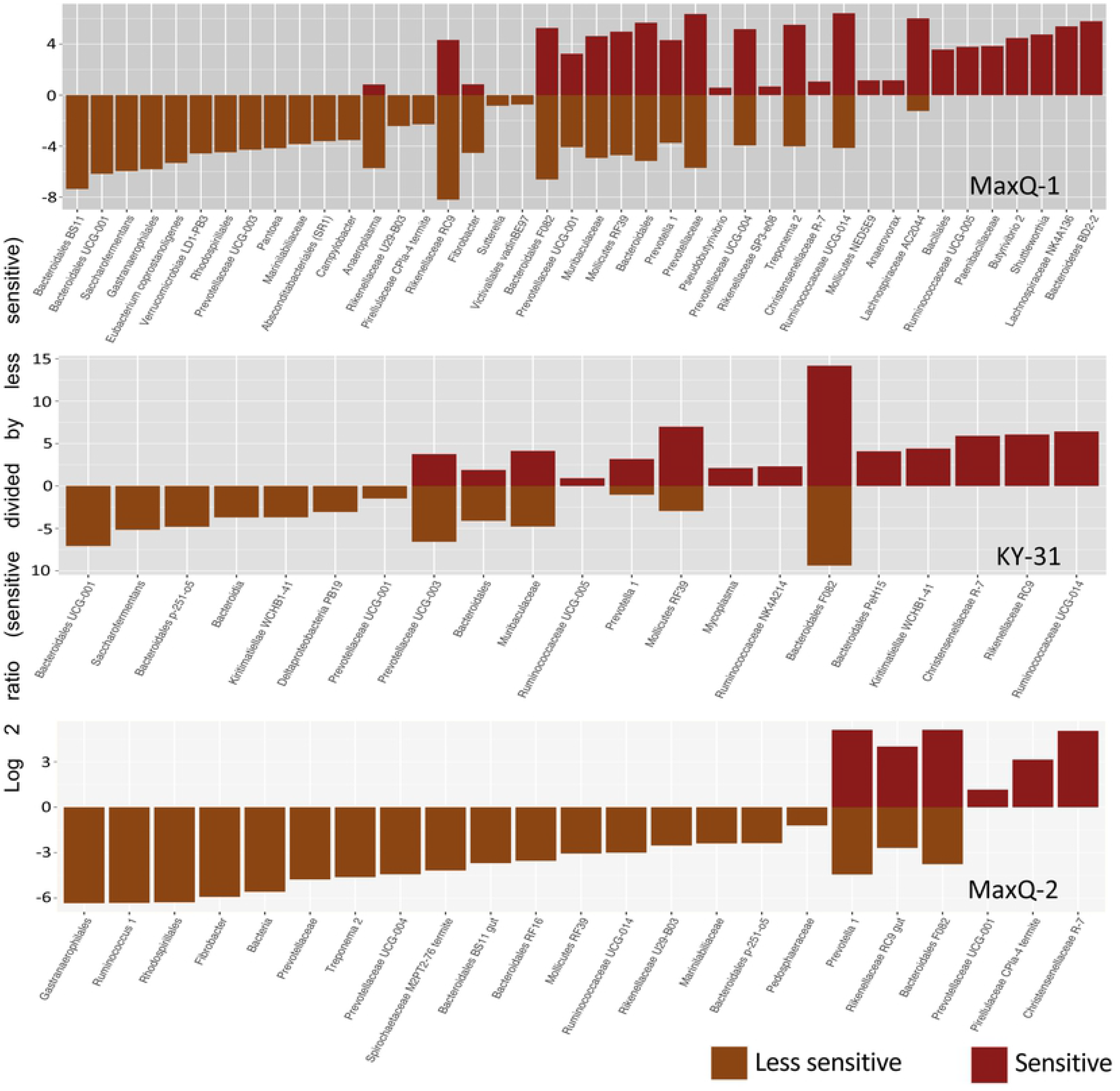
Archaea and bacteria apparently enriching in the rumen microbiome based on the sensitivity of the cattle to fescue toxicosis. Differentially abundant OTUs in sensitive with respect to less-sensitive steers under three grazing condition. Average Log2 values of the sensitive and less sensitive abundance ratio are shown on the Y-axis and the number of OTUs representing an assigned lowest taxonomic annotation are shown on the side of the respective bar.

The rumen microbiome of sensitive steers was associated with a higher abundance of Firmicutes. Especially, the species belonging to the R-7 group genera from the *Christensenellaceae* family showed a preferential association with the rumen of sensitive steers, being 2-, 59- and 32-fold more abundant during MaxQ-1, KY-31 and MaxQ-2 grazing, respectively, in this set compared to the less sensitive animals (Fig. 9). In addition to elevated Firmicutes abundance, a higher association of certain members of the Tenericutes family, specifically the Mollicutes order (Mollicutes RF39, *Mycoplasma*, and Mollicutes NED5E9) was observed for the rumen of sensitive steers.

## Discussion

The previous studies on the rumen microbiome of cattle consuming Tall Fescue have primarily focused on the possibilities and mechanisms of microbial degradation of ergovaline as produced by the fungal endophyte of the KY-31 cultivar (11, 17–19, 46, 47). In some of these cases the fecal microbiome was employed as a proxy (17, 18, 47). The current study had the goals to determine whether the toxic and non-toxic cultivars establish unique types of rumen microbiomes and if the cattle’s relative sensitivities to Tall Fescue Toxicosis influence the microbiome characteristics. Considering that the fungal endophyte of KY-31 produces ergovaline and that of MaxQ does not, our experimental design called for the use of grazing of a KY-31 pasture as a perturbation in middle of MaxQ grazing, and the resultant changes in the composition of the rumen microbiome were recorded. This effort had two major outcomes. One concerned a methodological aspect. It was observed that an analysis that targeted the sessile and planktonic components separately uncovered more changes in the microbiome than the one that was focused on the overall microbiome. Thus, an effective study of rumen microbiome metabolism cannot rely on the analysis of rumen fluid or the whole microbiome only (19, 48, 49). The other was the finding that KY-31 was a dominant remodeler of the microbiome composition. A return from KY-31 to MaxQ grazing did not restore some of the original MaxQ-type features. One of these cases was a severe reduction in the abundances of OTUs assigned to certain key rumen cellulolytic bacteria. The location within the microbiome, sessile or planktonic segment, where Firmicutes and Bacteroidetes were found in high abundances, changed when cattle were moved from MaxQ to KY-31 grazing, and the original status was restored upon return to MaxQ. *As elaborated at the end of the Discussion,* these remodeling events likely represent alterations in the microbial team composition and/or task allocations while preserving a broader ruminal metabolic function. *The more intriguing aspect of these observations* was that some of the KY-31 instituted changes lingered even after a switch to MaxQ grazing. This represents a major departure from what is seen with various perturbations in the ruminal conditions. For example, for bloat or acidosis upon the restoration of the normal ruminal pH the microbiome returns to the pre-disturbance stage (50, 51). There were indications that KY-31 grazing led to the enrichment of bacteria that likely possessed capabilities of degrading ergot alkaloids. The cattle’s relative sensitivities to fescue toxicosis had limited effect on rumen microbiome composition. The KY-31 grazing seemed to generate a rumen microbiome with lowered methane production potential. All these observations were significant, as the three-week long intervening KY-31 grazing, a terminal MaxQ grazing of similar duration, and much longer first MaxQ period offered sufficient times for establishing stable microbiome compositions; in cattle the sign of fescue toxicosis appears in 5-15 days of consuming toxic tall fescue (5) and the composition of fecal bacterial community alters within 14-28 days of a shift from a mixed ration to MaxQ or KY-31 (17). The rumen microbiome compositions identified in this study were found to be distinct from fecal microbiomes of cattle that grazed tall fescue (17, 18).

At the phylum-level, the overall rumen microbiome of beef cattle grazing tall fescue MaxQ or KY-31 was most similar to those developed with Alfalfa than other diets (Table S6) (49–56). However, tall fescue provided 6-10 times more Tenericutes abundance and significantly lower level of Actinobacteria than Alfalfa (Table S6) (50). The MaxQ grazing, provided up to 2-fold higher abundance of Proteobacteria compared to our KY-31 set and all above mentioned cases (49–55) (Table S6). The rumen fractions (solid and liquid, representing sessile and planktonic microbial populations) and the grazing periods (MaxQ-1, KY-31, and MaxQ-2), but not the rumen depths influenced sample clustering (Fig. 2 and Table 1). The fescue toxicosis sensitivity of the animals was significantly correlated with only one effect (Fig. 3): the rumen of less-sensitive cattle contained slightly higher abundances of Bacteroidetes and Fibrobacteres while that of sensitive cattle carried a bit more Firmicutes, especially *Christensenellaceae* family members, and Tenericutes (Fig. 9).

Our analyses focused on the rumen fractions identified a potentially major microbiome remodeling event. The MaxQ1 to KY-31 shift reduced the species diversity of the fiber-attached population and increased that of the liquid-associated population, and it caused an opposite effect for MaxQ-1 and MaxQ-2 grazing (Fig. 1). A phylum level observation was even more dramatic (Fig. 4A). A move from MaxQ to KY-31 grazing reduced the relative abundance of Firmicutes and elevated that of Bacteroidetes in the sessile fraction and the opposite change occurring in the planktonic fraction, and a return to MaxQ brought back the original composition, with a shift to a higher abundance of Firmicutes and a lowered level for Bacteroidetes in the sessile fraction and the reverse for the planktonic fraction. These effects were not seen at the total microbiome level as both KY-31 to MaxQ provided similar overall abundances of Firmicutes and Bacteroidetes in the rumen, which in combination constituted >80% of the bacterial population. We envisage two reasons for the remodeling event. One would be the toxicity of the ergot alkaloids released from the fibers in the solid fraction and the other the availability of these compounds as high value nutrients in the liquid fraction. However, the reversibility as observed for the sessile and planktonic Firmicutes and Bacteroidetes did not hold for certain features that are key to an optimal functioning of the rumen.

For the sessile fraction of the rumen microbiome, the transfer of cattle from MaxQ to KY-31 grazing caused an increase in the abundance of the Bacteroidetes, such as saccharolytic *Prevotella 1* and *Paraprevotella* species and RC9 of the *Rikenellaceae* family, and lowered the abundance of the Firmicutes representing major rumen cellulolytic populations with members such as *Pseudobutyrivibrio* and *Butyrivibrio* 2 (Fig. 4B and Fig. 7). The opposite was the case for the planktonic population. There was also a decrease in the abundance of fibrolytic *Fibrobacter* species in the solid fraction (Fig. 6). This void in the sessile population was filled with an enrichment of cellulolytic firmicutes belonging to the *Ruminococcaceae* and *Lachnospiraceae* families (Figs. 4B and Fig. 7), and the Eubacterium genus (*E. coprostanoligenes*, *E. ventriosum,* and *E. ruminantium*). The abundance of *E. ruminantium*, a typical rumen cellulolytic bacterium (57), was elevated substantially during KY-31 grazing and remained so in MaxQ-2. Bloat and acidosis are known to affect *Fibrobacter succinogenes* and the Firmicutes species such as *Ruminococcus albus* and *Ruminococcus flavefaciens* (50, 51, 58). Acidosis is associated with pH values of 5.0-5.6 (51, 59), whereas under our grazing conditions, the rumen pH remained constant (7.00 ± 0.24). Thus, certain factor(s) associated with the KY-31, such as ergovaline, were possibly deleterious to *Fibrobacter, Pseudobutyrivibrio* and *Butyrivibrio* and this loss was compensated by select Firmicutes.

For the planktonic segment of the microbiome, KY-31 grazing not only altered the level of the cellulolytic and non-cellulolytic carbohydrate hydrolyzing bacterial population but also brought novel possibilities for organic and cellular material degradation activities. For example, the increased abundance of planktonic Firmicutes likely not only strengthened the ruminal cellulolytic activity and compensated for the above-mentioned loss in the sessile fraction, but also enhanced the ability to degrade plant derived organics. A member of this phylum, a *Coprococcus* species, was present at an enhanced level during KY-31, and these organisms degrade plant toxins such as nitropropionic acid, nitropropanol, and phloroglucinol (60–62). Similarly, the *Anaeroplasma* and *Mycoplasma* species as well as the bacteria from the RF9 family of the Mollicutes order, all of which belong to the Tenericutes phylum and were present in elevated abundances during the KY-31 grazing, would have provided two benefits. First, their saccharolytic and proteolytic activities likely compensated for the lowered Bacteroidetes abundance in the planktonic community. Second, their bacteriolytic and proteolytic activities were potentially useful in recycling dead cells (63). In this context it is noted that not all rumen mycoplasmas are bacteriolytic (63). Proliferation of acetogens such as *Acetitomaculum*, sulfate-reducing *Desulfovibrio*, species, and propionate-producing bacteria of the *Rikenellaceae* and *Erysipelotrichaceae* families indicated that under KY-31 grazing the contribution of methanogenesis as a hydrogen sink might have been lowered (64). This suggestion is consistent with the observation that during KY-31 grazing the rumen microbiome carried *Methanosarcina*, which are capable of living without hydrogen (65, 66), in a moderately higher abundance than that with MaxQ-1 and MaxQ-2 (Fig. 5). Also, since the methanogen abundance did not experience a major change, KY-31 or its degradation products were not likely toxic to these archaea.

The transition of the cattle from KY-31 to MaxQ provided a rather unexpected outcome. Even after 23 days of MaxQ-2 grazing following the KY-31 period, the rumen microbiome of the cattle did not return completely to the MaxQ-1 stage (Fig. 2 and 8), and exhibited some of the KY-31 stage characteristics (Fig. 2 and 4). This observation contrasted those made in studies on acidosis (51) or involving an administration of butyrate into the rumen (67), where the microbiome fully regains the pre-disturbance characteristics within 7 days of the termination of the disturbance (51, 67). The core ruminal microbiome during the MaxQ-2 had the highest abundance of Firmicutes, up to 50% of the total sequence, in both solid and liquid fractions (Fig. 4A, Table S2). One of the special features of this period was the higher abundance of Spirochaetes, specifically the species of the *Treponema 2* genus, and organisms belonging to the *Pedosphaeraceae* family of Verrucomicrobia phylum (Fig. 4). Several organisms such as the Firmicutes of the *Tyzzerella*, *Ethanoligenes* and *Succiniclasticum* genera as well as those belonging to *Bacteroidales RF16* order and *Rikenellaceae* RC9 families of the Bacteroidetes phyla that were present in the MaxQ-1 rumen microbiome in high abundance and were affected due to KY-31 grazing were able to re-establish substantial presence in the rumen under MaxQ-2 grazing. However, the abundances of *Fibrobacter* and *Butyrivibrio* 2 species in the sessile fraction, which were severely reduced in the KY-31 period, did not return to the MaxQ-1 type level during MaxQ-2 grazing (Fig. 8). On the other hand, the *Treponema* species which were present at high levels during the MaxQ-1 grazing and had a reduced presence at the KY-31 phase, gained significant abundance in the sessile fraction with the MaxQ-2 (Fig. 7). These bacteria are known to improve the cellulolytic function of cellulose degraders, especially *Fibrobacter* (68–70), and this property is consistent with the significant presence of *Treponema* during MaxQ-1 grazing where *Fibrobacter* species were present in high abundance. For MaxQ-2, where *Fibrobacter* abundance was low, *Treponema* species likely assisted other cellulolytic organisms.

There have been efforts to use the fecal microbiome as a proxy for monitoring the activities in the rumen under particular diets (49, 71) and ruminal conditions (50), and to identify particular fecal microbes as biomarkers for fescue toxicosis (17, 18, 47). We compared our findings with the published information on the fecal microbiome of cattle that grazed tall fescue. In two studies where cattle were moved from a mixed ration to MaxQ and KY-31 in parallel (17, 18) or to KY-31 only (47), *Coprococcus 1* and several other Firmicutes belonging to the *Ruminococcaceae* and *Lachnospiraceae* families, *Paraprevotella* species from Bacteroidetes phylum, and Tenericutes of *Mycoplasmataceae* family were found in higher abundances in the feces of KY-31 tall fescue-fed cattle (17, 18, 47). Similar changes were observed in our MaxQ-1 to KY-31 shift (Fig. 7). The fecal microbiome of cattle grazing tall fescue also contains Proteobacteria, Euryarchaeota, and Lentisphaera phyla at abundances similar to those observed by us for the rumen (17, 18, 47). However, unlike in the latter, in a fecal microbiome Actinobacteria and Firmicutes are found in significantly higher abundances, while Bacteroidetes, Fibrobacteres, and Tenericutes are underrepresented (17, 18, 47). Similar observations have been made with alfalfa (50) and a mix of bermudagrass and Dallisgrass (49) (Table S6). Such differences are expected as the foregut deals with heterogenous plant materials and in the hindgut undigested plant fibers are degraded (49, 72–74). Also, the hindgut selects for microbes that can tolerate bile salts (73), which may inhibit many Gram-negative bacteria such as Bacteroidetes and Fibrobacteres and help to enrich Gram-positive bacteria such as Actinobacteria and Firmicutes (73). In Holstein steer, the foregut and hindgut microbiomes share only 30% of the species (17, 20). Thus, in many aspects a fecal microbiome of cattle grazing tall fescue does not represent that of the corresponding rumen.

The results of an *in vitro* enrichment study suggest that certain *Clostridiaceae* and *Prevotellaceae* can degrade ergovaline by leveraging their amino acid degradation machineries (46). From an analysis of the fecal microbiomes of cattle grazing toxic and nontoxic tall fescue, certain *Paraprevotella* and *Coprococcus 1* species and several members of the *Ruminococcaceae*, *Lachnospiraceae* and *Mycoplasmataceae* families have been identified as potential ergovline degrader (17, 18, 47). A study on the planktonic segment of the rumen microbiome of pregnant ewes have raised similar possibilities for certain bacteria belonging to the Lachnospiraceae and Veillonellaceae families of the Firmicutes phylum and Coriobacteriaceae family of Actinobacteria phylum (19). From the inferred metabolic capabilities of the bacteria that were enriched in the rumen as the cattle shifted from MaxQ-1 to KY-31 grazing, we have identified *Coprococcus 1, Paraprevotella*, *Prevotella 1*, *RC9*, *Mycoplasma*, *Anaeroplasma* and several members of the *Ruminococcaceae* and *Lachnospiraceae* families as potential degraders of ergovaline. These similarities between the hypotheses arising from rumen and fecal microbiome studies (current report and (17–19, 47)) indicate that ergovaline metabolism in the rumen and large-intestine occur through similar microbial processes. Ergovaline generated from KY-31 in the rumen is known to remain available in the large intestine of cattle (16, 75).

We envisage two mechanisms that drove the observed KY-31-instigated remodeling event. In one ergovaline was the remodeling agent. The other considers that KY-31 is more than simply being MaxQ with ergovaline, and therefore, these two forages selected for two microbiomes or microbial teams of distinct compositions. Clearly, with MaxQ more Firmicutes were present in sessile forms or as fiber adhered degraders than in the free forms, and the majority of the Bacteroidetes were located in the planktonic segment, and KY-31 selected for an opposite distribution. Similarly, MaxQ helped to enrich a cellulose metabolizing community made up of *Fibrobacter*, *Pseudobutyrivibrio* and *Butyrivibrio* 2, and for the same overall function a consortium of *Eubacterium* species and certain members of the *Ruminococcaceae* and *Lachnospiraceae* families was engaged under KY-31 grazing. Distinct team configurations were employed for the saccharolytic functions as well. These are simpler explanations and the systems certainly were comprised of more complex compositions and metabolic networks. The observations such as there was an enhanced need for Tenericutes for the degradation of KY-31 and some of the KY-31 specific features were retained during the MaxQ grazing period that followed were parts of these complexities. Often, beef cattle are raised by grazing tall fescue until the age of 6-12 months, and then, the animals are placed on feedlots until the age of 18-21 months for finishing (26). It is now clear that the cattle groups that graze KY-31 and MaxQ will arrive on feedlots with distinct microbiomes, and the characteristics developed with KY-31 will likely linger. Thus, a fuller understanding of the metabolic details of the complex systems described above will help to develop strategies not only for dampening or eliminating the fescue toxicosis but also for improving the feed utilization efficiencies at both the Tall Fescue grazing and finishing step. It will also reveal new biomass degradation systems of applies value. One possible next step towards obtaining more granular views of the identified remodeling event and the metabolic potentials of the rumen microbes involved would be the generation and analysis of metagenome assembled genomes (MAGs). This step could be followed by transcript, protein and metabolite level analysis.

## Acknowledgements

We thank Danny Eanes for expert help in the construction of the sampling device, Sierra R. Guynn and Terry Swecker for the cannulation of the steers, Brad Ellis, Lauren Rush and Endang Purwantini for assistance in sample collection, and Brian Hedlund and Shrikant Bhute for advice of obtaining 16SrRNA-V4 sequencing and introducing BHK to QIIME.

## Supporting information

**Method S1.** Rumen sample collection

**Method S2.** DNA extraction

**Fig. S1.** Custom made sample collection system

**Fig. S2.** Sample rarefaction of tall fescue rumen microbiome

**Table S1.** Rumen samples included and excluded from the analysis

**Table S2.** Core microbiome - OTUs present with 95% Prevalence

**Table S3.** Microbiota shift due to grazing transitions

**Table S4.** Microbiota shift in specific rumen fraction (solid and liquid) due to grazing transitions

**Table S5.** Differentially abundant prokaryotic OTUs in the rumen of sensitive steers with respect to that of less sensitive steers

**Table S6.** Phylum-level microbiota comparison between steers graze on two tall fescue cultivars and other diets

